# JRSeek: Artificial Intelligence Meets Jelly Roll Fold Classification in Viruses

**DOI:** 10.1101/2025.01.27.635132

**Authors:** Jason E. Sanchez, Wenhan Guo, Chunqiang Li, Lin Li, Chuan Xiao

## Abstract

The jelly roll (JR) fold is the most common structural motif found in the capsid and nucleocapsid of viruses. Its pervasiveness across many different viral families motives developing a tool to predict its presence from a sequence. In the current work, logistic regression (LR) models trained on six different large language model (LLM) embeddings exhibited over 95% accuracy in differentiating JR from non-JR sequences. The dataset used for training and testing included sequences from single JR viruses, non-JR viruses, and non-virus immunoglobulin-like β-sandwich (IGLBS) proteins which closely resemble the JR fold in structure. The high accuracy is particularly remarkable given the low sequence similarity across viral families and the balanced nature of the dataset. Also, the accuracy of the models was independent of LLM embeddings, suggesting that peak accuracy for predicting viral JR folds hinges more on the data quality and quantity rather than on the specific mathematical models used. Given that many viral capsid and nucleocapsid structures have yet to be resolved, using sequence-based LLMs is a promising strategy that can readily be applied to available data. Principal Component Analysis of the Bert-U100 embeddings demonstrates that most IGLBS sequences and a subset of JR and non-JR sequences are distinguishable even before the application of the LR model, but the LR model is necessary to differentiate a subset of more ambiguous sequences. When applied to double JR folds, the Bert-U100 model was able to assign the JR motif for some viral families, providing evidence for the model’s generalizability. However, for other families, this generalizability was not observed, motivating a future need to develop other models informed by double JR folds. Lastly, the Bert-U100 model was also able to predict whether sequences from a dataset of unclassified viruses produce the JR fold. Two examples are given and the JR predictions are corroborated by AlphaFold3. Altogether, this work demonstrates that JR folds can, in principle, be predicted from their sequences.

## Introduction

The single jelly roll (SJR) is the most common type of large-scale structural motif found in viral capsid proteins (CPs) and nucleocapsid proteins (NCPs), representing approximately 28% of entries in an analysis by Krupovic and Koonin.^1^ Remarkably, the SJR fold is found in all domains of life and is involved with a wide array of functions such as carbohydrate-binding, protein stabilization, and nucleosome assembly, to name a few.^2–7^ For many viral families, the SJR is the principal structural domain necessary for proper viral assembly. It is postulated this function arose from diversification events acting on cellular (non-viral) SJR proteins before the emergence of the last universal cellular ancestor.^1^ One line of evidence for this claim is that soluble tumor necrosis factor and Apo-L-related-leucocyte-expressed-ligand-1 (sTALL-1)—an extant cellular TNF-like protein containing a SJR domain—can form 60-subuint aggregates that resemble viral particles.^8^

In light of this finding, researchers propose that viral SJR folds may have evolved to recruit one another—a necessary step for the production of nascent viral particles. The ability of SJR proteins to bind the viral genome, another requirement, could have come about by augmenting the core SJR structure. In fact, for many ssRNA and ssDNA viruses, there are large, positively-charged, unstructured N-terminal domains which serve to interact with the viral genome.^9,10^ Indeed, viral jelly roll (JR) assembly is an active area of research, and many mechanisms have been proposed to explain how JR proteins recruit one another to form mature viral particles.^11^

The adaptability and ubiquity of SJR folds and their abundance among certain viral CPs, motivates a desire to predict these from a given sequence. This is challenging as many structurally similar viruses have little recognizable sequence similarity. For instance, when the structure of the southern bean mosaic virus (SMBV) was elucidated, it was shown to bear a striking resemblance to that of the tomato bushy stunt virus (TBSV). These two viruses—the first icosahedral viruses to have their structure uncovered— were later determined to share a JR fold in spite of sharing little sequence similarity.^12–14^ Before attempting to understand how two very different sequences like these could result in a similar structure, it is necessary to explore the JR structure itself.

The SJR fold, also known as the wedge-shaped fold, RNA capsid β-barrel or β-sandwich, is formed by eight antiparallel β-strands curved to form an incomplete β-barrel-like or β-sandwich structure.^15–17^ A variation of the more common Greek key structure, the SJR fold is part of the larger family of protein structural motifs whose main feature is the preponderance of β-strands.^18^ In SJR folds, these β-strands are lettered B through I along the path formed by the peptide. An example SJR fold from the SMBV virus and the larger viral particle formed by SJR CPs are shown in Figures 1A and 1B. Notably, SJR folds exhibit a wide range of length spanning between 180 amino acids on the low end to 580 amino acids, with much of this size difference attributed to the disordered loops connecting the β-strands.^15,19–21^

**Figure 1.**
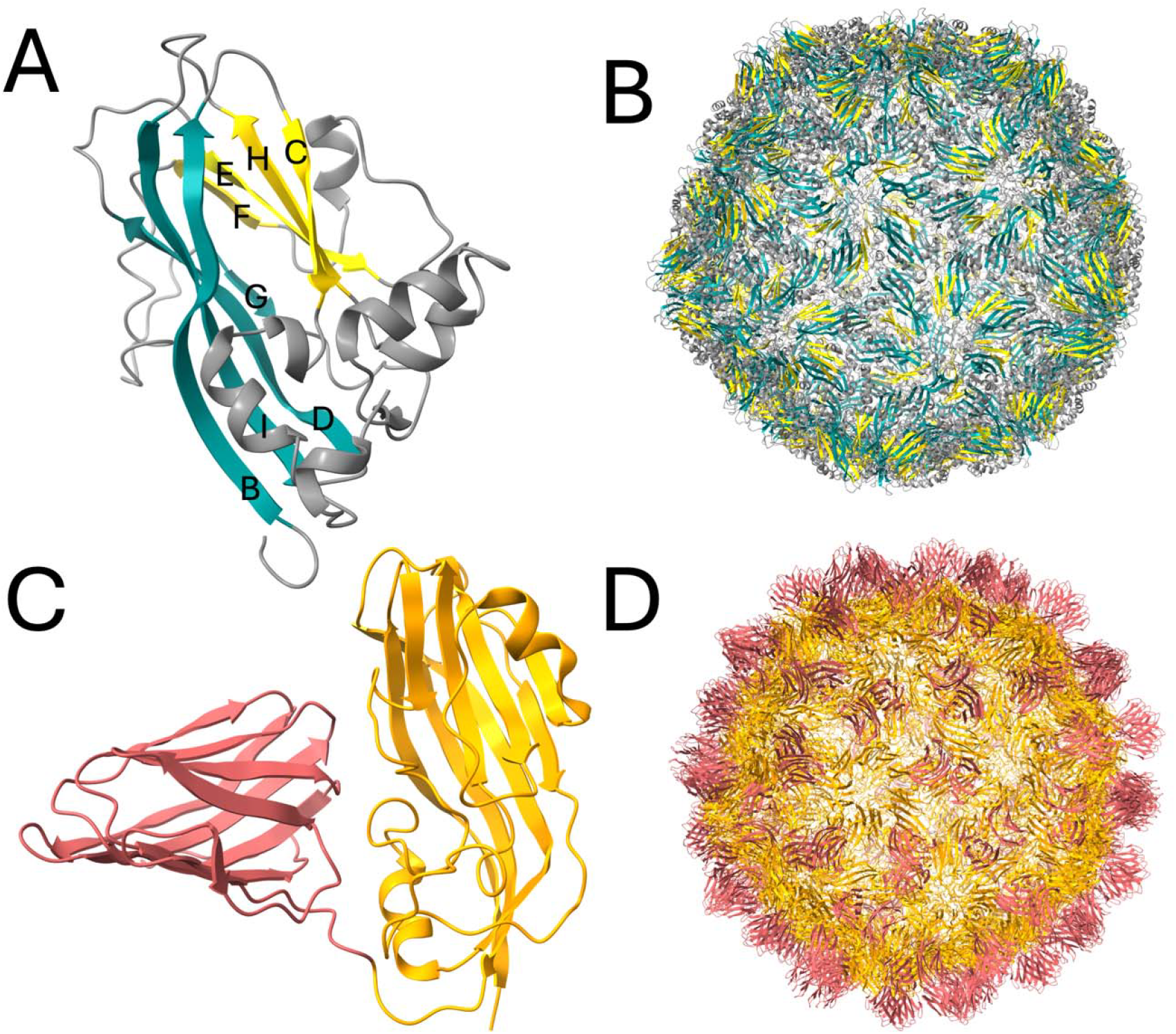
Example SJR and DJR folds from the CPs of the SBMV (PDBID: 4SBV) and TBSV (PDBID: 2TBV) viruses.^22,23^ A) The two β-sheets from the SJR fold are colored teal and yellow. The lettering scheme from B to I follows the length of the peptide chain starting from the N-terminus. B) The icosahedral SBMV is made up of repeated CP bearing the SJR folds. C) The DJR fold is composed of two adjacent SJRs, here colored orange and pink. D) The icosahedral TBSV is made up of repeated CPs bearing the DJR fold.

In addition to the SJR, there also exists a double jelly roll (DJR) motif. As the name suggests, the DJR is made up of two consecutive SJRs. Figures 1C and 1D depict a DJR fold from the TBSV along with its associated virion. It is proposed that the DJR fold might have come about by gene duplication.^1,24,25^ This is supported by viruses of the Sphaerolipoviridae family which encode two CPs each with a SJR fold. ^26–28^ It is possible that these separate SJR folds might have merged in an ancestor of modern day DJR viruses. In terms of a sequence-based effort to predict SJRs, DJR present a unique opportunity. Presumably, a method which detects SJRs would detect DJRs making for an external test-case for a sequenced-based method. A tool which can predict SJRs ought to generalize to some DJRs in spite of the two folds coming from different viral families.

With these facts in mind, a natural choice for realizing sequence-based predictions is artificial intelligence (AI). Specifically, Large Language Models (LLMs)— which can extract meaningful context from ordered text— have already been used for many biological applications and are well-positioned to make predictions about the JR content of proteins.^29–31^ Not limited to human language, LLMs can make sense of many problems which can be understood by a symbolic representation. Of course, in the field of protein biology, peptide sequences are a language of sorts that encode information about the one-dimensional structure of proteins.

Perhaps one of the most important use cases for protein-focused LM is 3D structure prediction. Among the most well-known tools for this purpose are AlphaFold, Evolutionary Scale Modeling Fold (ESMFold), and RoseTTAFold.^32–34^ Given only a sequence, these models return atomic-level predictions for the 3D structure proteins at impressive speed and accuracy. Although these tools represent a tremendous breakthrough in understanding how a sequences might fold into functional proteins, interpreting the output of these at scale can be time-consuming. Indeed, though protein structure prediction is well-suited to uncover JR folds, a visual inspection or computational analysis of the output is required to confirm the result. To circumvent this, a sequence-only approach is proposed in this work.

Here we describe other examples of LLMs that have been applied successfully for different sequence-based tasks. A review by Savojardo et al. compiles examples of efforts aimed at predicting structural motifs.^35^ Those tools which use LLMs can be grouped into three main categories: transmembrane protein prediction, subcellular localization prediction, and signal peptide detection. The use case most closely aligned with this study is transmembrane protein prediction. This is because transmembrane proteins are either ⍰-helix-based or β-barrel-based—a structure resembling JR folds.^36–39^ Among the tools listed, TMbed is able to accurately predict whether a sequence results in a transmembrane or non-transmembrane protein with an accuracy of 94% for β-barrel and 98% for ⍰-helical proteins.^36^ This is particularly remarkable given that β-barrel proteins were far outnumbered by their ⍰-helical counterparts (65 versus 593). Owing to the similarity of JR folds to β-barrels, it is natural to explore whether a similar approach can be used to identify the JR motif.

Underpinning TMbed is ProtT5, a LLM trained on the Uniref50 dataset of protein sequences.^40,41^ In short, ProtT5 uses the T5 LM architecture but in place of human language, amino acid sequences from all domains of life are used.^40,42^ T5 and various other LLMs like it make use of a concept called “transfer learning” where a large body of input (e.g. English or protein sequences) is used to train a LLM to recognize patterns in the text. Often, this training occurs by an unsupervised fashion whereby certain words—or, by analogy, amino acid sequences—are “masked” and the model is tasked with predicting the actual value of the masked words. In this way, the model learns the underlying “grammar” of the language without needing to be explicitly trained. After this pre-training stage, a model like T5 can produce a vector-representation of every word (amino acid) in a sequence. These so-called embeddings encode information about the word itself as well as the context around it (i.e. future and past words). In this way, embeddings are primed to learn about emergent aspects of a protein (such as secondary structure) and may be used for further downstream tasks that require fine-tuning for a specific purpose—such as transmembrane motif prediction.

The creators of ProtT5, also made available other LLMs pre-trained on Uniref50, Uniref100, and the Big Friendly Dataset (BFD) in a repository entitled ProtTrans. ^40,41,43^ The LLMs used in the repository were T5, Electra, BERT, Albert, Transformer-XL, and XLNet. ^42,44– 48^ Importantly, these models make use of the transformer architecture first described by Google in 2017.^49^ The transformer helped remedy two problems that had plagued many LLMs at the time: long-distance dependence between words in a sentence and difficulty in parallelizing learning.^49,50^ The transformer architecture, which uses a technique called self-attention, was able to address these two issues, making the models more accurate on longer sequences and shortening the time required for training.

In this study, previously trained transformer models from ProtTrans are leveraged to make predictions about the presence of JR folds from a dataset of JR proteins and non-JR proteins. The non-JR sequences in this study were from viral and immunoglobulin-like β-sandwich (IGLBS) sequences. Immunoglobulin-like proteins were chosen because of their structural similarity to JR proteins. Surprisingly, at the fine-tuning stage of transfer learning, a logistic regression (LR) function is all that is needed to achieve remarkable performance for the embeddings used. The models were compered in terms of their accuracy and speed. The embeddings of the top performing model, Bert-U100, were analyzed in greater detail.

Additionally, the Bert-U100 model was used to predict the presence of JR folds in a dataset comprised of DJR sequences and β-strand-dominated proteins. Lastly, novel JR folds were predicted from a set of unclassified viruses and their structures were confirmed with AlphaFold3. The LR models are made available at: https://github.com/LiLabBioPhysics/JRSeek.

## Results and Discussions

### Dataset Composition and Sequence Analysis

Representative structures of all motifs used is this study are shown in Figure 2. The characteristic eight antiparallel β-sheet sandwich for SJRs and the structurally similar IGLBS are shown on the top row. All motifs which are not SJR, DJR, or IGLBS are referred to as viral non-jelly roll (VNJR). Notably, no VNJR motif bears a resemblance to the SJR or IGLBS motifs. For instance, the most salient characteristic of five VNJR motifs (Alpha Helical, Arena-like, Borna-like, Chymotrypsin-like, and Corona-like) is the dominance of ⍰-helical secondary structures. It stands to reason that a transformer model could predict the presence of JRs by assessing putative ⍰-helical versus β-sheet content. Even the remining four motifs—though possessing a greater proportion of β-sheet residues—would likely not challenge a model. None of these (Beta Sheet, Chymotrypsin-like, HK-97-like, Corona-like, and Reo-like) closely emulate the rolled β-sheet sandwich characteristic of JRs. The Chymotrypsin-like motif, which most closely resembles the SJR in that it contains an interface separating two adjacent sets of β-sheets, has a distinct concave-like (not barrel-like) shape.

**Figure 2.**
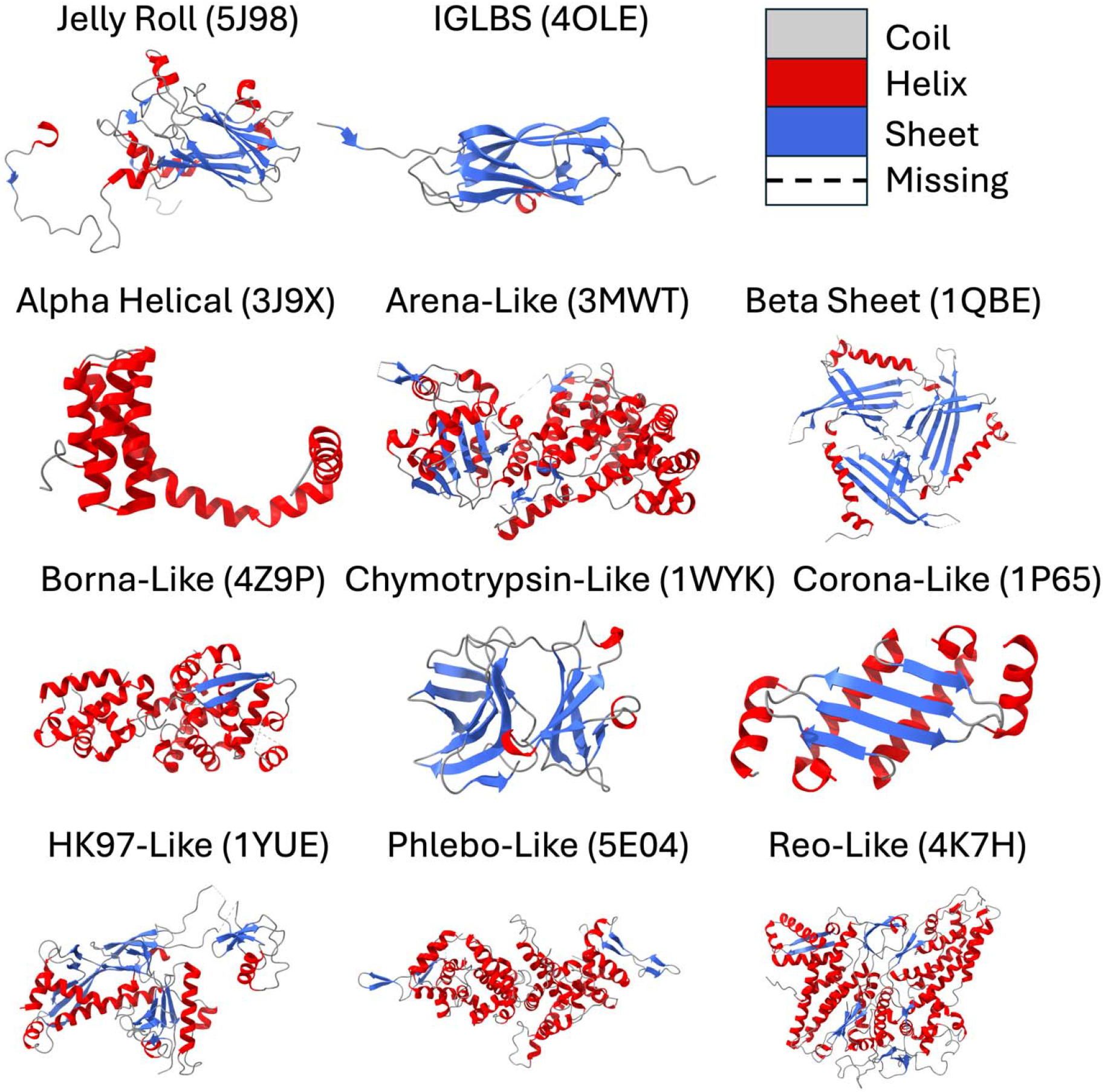
Examples of motifs present in the study. The structures are color coded to show coil (grey), ⍰-helical (red), and β-sheet (blue) regions of proteins. PDB IDs are shown in paratheses.

A similarity analysis of sequences of the top twenty most populated viral families (Figure 3A) reveals low-to-moderate (between 0% and 50%) average sequence identity for datapoints within the same viral family (diagonal entries), though some families (e.g. Arteriviridae) exhibit more intra-family similarity. Because a robust model is ideally exposed to a diverse set of sequences, having low average identity represents challenging training-conditions. More varied sequences allows a model to produce more credible predictions over a larger set inputs. Specifically, for the task at hand, low intra-family sequence identity suggests that sequences within each family come from many different genera and/or species and bolster the predictive capacity of the model.

**Figure 3.**
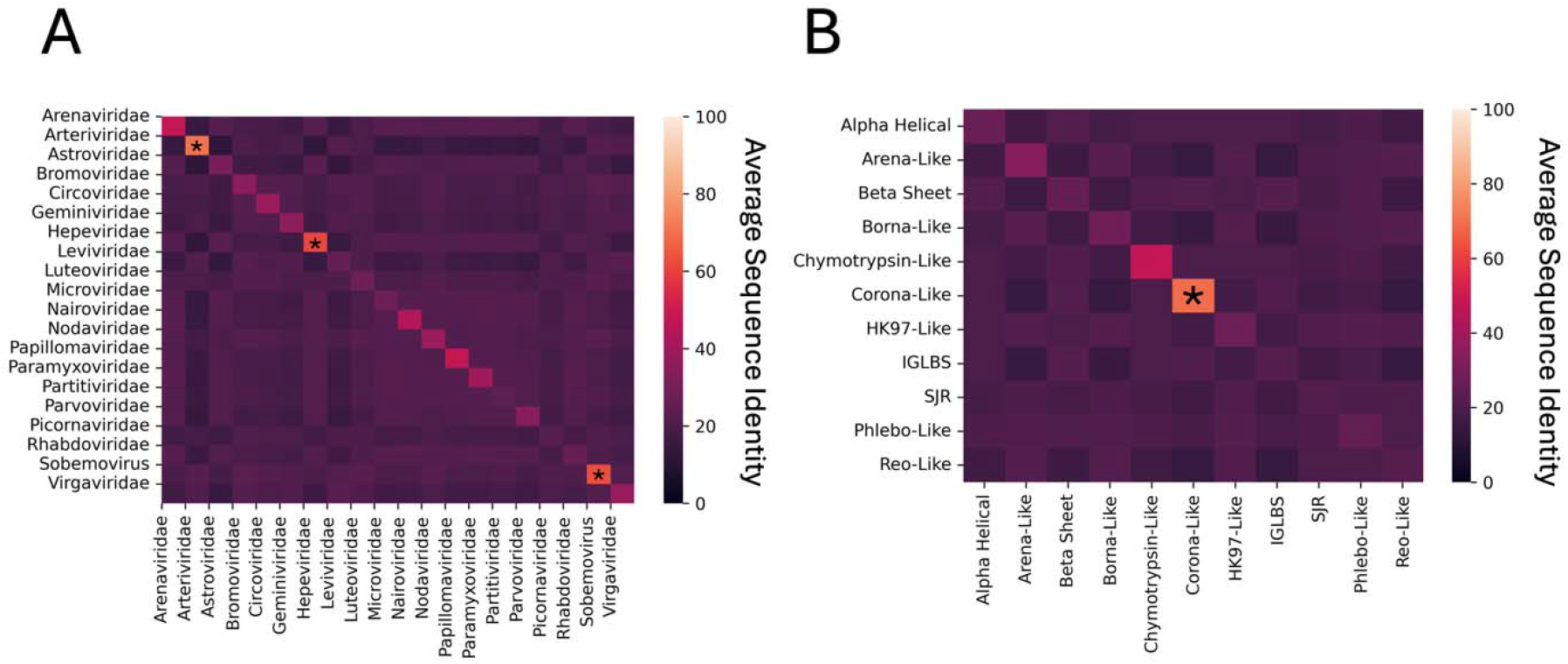
Average sequence identity heatmaps. Asterisks along the diagonal denote entries with > 50% average sequence identity A) Among the top 20 families with the most sequences, most exhibit low intra-family and inter-family sequence identity. B) A per-motif sequence identity analysis reveals little identity between sequences.

Focusing on inter-family sequence identity, large swaths of purple off-diagonal boxes in Figure 3A confirm that the viral families used in this work are genetically diverse. Further, visualization of sequence identities by motif (Figure 3B) reveals little average sequence similarity per motif. The one motif that does exhibit a moderate amount of sequence identity is the Corona-like motif. This can be explained by the fact that all Corona-like NC sequences come from the Arteriviridae family of viruses which itself has high average inter-family sequence identity (Figure 3A). Lastly, the low sequence identity for SJR sequences, in particular, underscores that a method which aims to predict SJRs from sequences may not solely rely on similarly/homology from one type of SJR.

### Model Performance

Following data cleaning, six LLMs were used to featurize the sequences. These were the Bert-BFD, Bert-U100, T5-BFD, T5-U50, XLNet-U100 and Albert-U100 models. Embeddings were used to train and test six separate LR models via a 70/30 train test split. The models were compared to evaluate whether the capacity to discriminate JR sequences is embedding-dependent. Relevant performance metrics used for model selection are shown in Table 1.

**Table 1.** Performance Metrics.

| Model | Albert-U100 | Bert-BFD | Bert-U100 | T5-BFD | T5-U50 | XLNet-U100 |
| --- | --- | --- | --- | --- | --- | --- |
| Accuracy | 0.98 | 0.97 | 0.97 | 0.97 | 0.99 | 0.95 |
| ROC AUC | 1.0 | 0.99 | 0.99 | 1.0 | 1.0 | 0.99 |
| False Positive Rate | 0.02 | 0.03 | 0.02 | 0.03 | 0.01 | 0.09 |
| False Negative Rate | 0.02 | 0.02 | 0.03 | 0.02 | 0.02 | 0.03 |
| Balanced Set Embedding Time (minutes) | 48 | 9 | 9 | 29 | 29 | 24 |

The accuracy of all models at the classification threshold of 0.5 was ≥ 0.95, indicating all models able to correctly identify both JR and non-JR sequences. Accuracy is especially meaningful in this analysis as the dataset was balanced for JR and non-JR classes. The ROC AUC of Albert-U100, T5-BFD, and T5-U50 was approximately 1.0 and that of Bert-BFD, Bert-U100, and XLNet-U100 was 0.99 indicating performance was sustained at many classification thresholds. The ROC curve for the Bert-U100 model is show in Figure 4. The false positive rate (FPR) of all models was ≤ 0.03, with the exception of XLNet-U100 which had an FPR of 0.09— three times higher than that of the other models. For the false negative rate (FNR), all models had values ≤ 0.03. Taking all these metrics together, the data show five models did not meaningfully differ in their ability to discriminate JR sequences from non-JR sequence. As XLNet-U100 had a higher FPR, this model was ruled out for subsequent experimentation.

**Figure 4.**
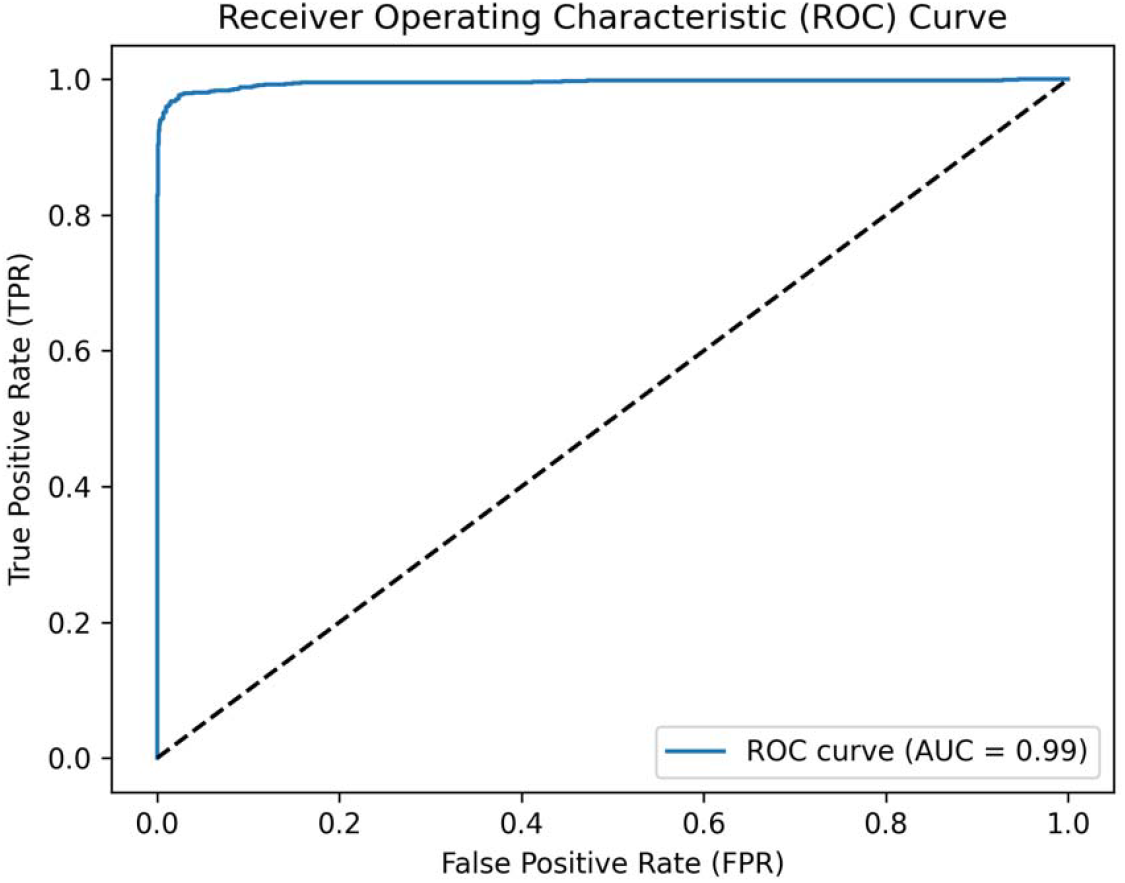
ROC curve for the test set using the Bert-U100 embeddings

Because the performance of the remaining models was very similar, speed was the next factor considered for model selection. The five remaining models could be grouped into three categories based on the time it took to develop embeddings (the most computationally demanding step). Albert-U100, though having the second-highest accuracy took about 48 minutes to produce embeddings. The next group is T5-BFD and T5-U50. T5-BFD underperformed T5-U50 in terms of Accuracy and FPR and both models took approximately 29 minutes to compute embeddings. Lastly, Bert-BFD and Bert-U100 performed almost identically with Bert-U100 having a more favorable FNR than Bert-BFD (0.02 versus 0.03) but having a less favorable FPR (0.03 versus 0.02). Both models took about 9 minutes to produce embeddings. On the basis of speed and model performance, either Bert-BFD or Bert-U100 could be chosen for subsequent analysis; however, Bert-U100 was selected for its lower FPR.

Having selected Bert-U100, Principal Component Analysis (PCA) of all sequences was conducted. To visualize the results, the transformed coordinates for all entries were plotted according to the first two principal components (Figure 5). Large regions of overlap in Figure 5A demonstrate diverse representation in the embeddings space in both the train and test sets. Three clusters with high proportions of SJR, VNJR, and IGLBS entries could be seen from the PCA (Figure 5B), which shows how the LR models learn to differentiate data from embeddings. The three ILBS, SJR, and VNJR clusters are located at the bottom, top, and right of the scatterplot, respectively. Also, there is a more ambiguous area between these clusters where ILBS, SJR, and VNJR sequences are all present.

**Figure 5.**
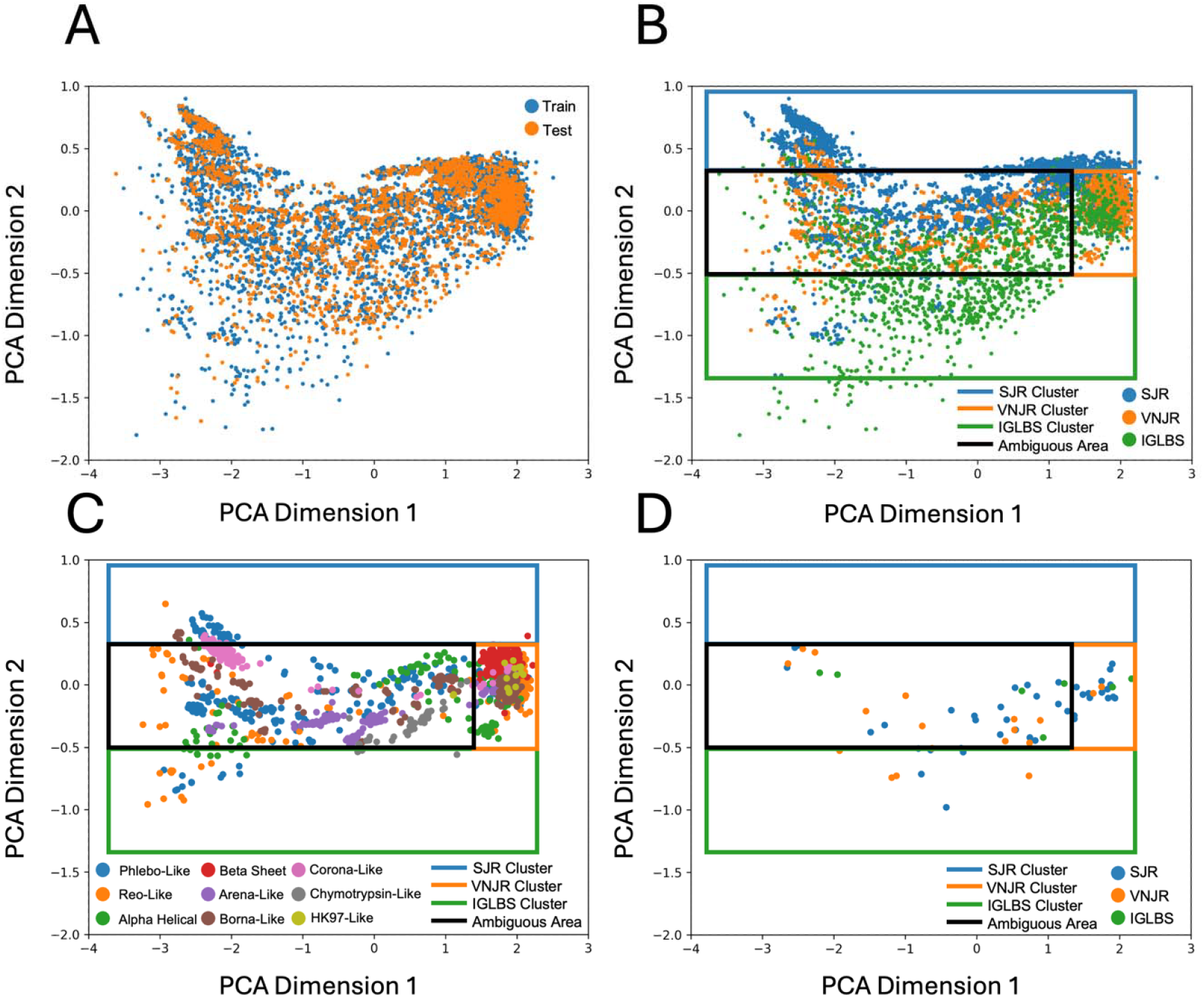
Scatterplots of Bert-U100 embeddings using the first two PCA dimensions. A) Datapoints in both the train and tests sets span a similar area, indicating the train/test split operation sampled embeddings evenly. B) Bert-U100 embeddings appear to capture differences in SJR, VNJR, and ILBS sequences. Three clusters, enclosed by blue, orange, and green rectangles—exhibit a high proportion of SJR, VNJR, and ILBS sequences, respectively. A black rectangle encloses an ambiguous area in the scatterplot where SJR, VNJR, and ILBS labels datapoints highly shuffled. C) Among the VNJR sequences, those with the same motif tend to cluster together. The Beta Sheet and the HK97 motifs are found predominantly in the VNJR cluster. D) Misclassified datapoints from the test set are shown. Many misclassified points are in the ambiguous area between clusters.

Firstly, the ILBS cluster is large and disperse. This expanse could be explained because of the origins of the ILBS data. Data from the SCOP dataset comes from eukaryotic, bacterial, archaeal and viral sequences, implying that ILBS sequences are diverse and span a large imbedding space. This contrasts with the SJR and VNJR sequences from NCBI which are solely from viral sources. Previous findings reported by the creators of the ProtTrans models show similar results. In the previous work, the T5-U50 models were shown to differentiate between viral, bacterial, archaeal, and eukaryotic sequences after t-SNE clustering.^40^ This result explains why the LR models in the current work are able to readily distinguish ILBS from SJR folds in spite of their structural similarities. Interestingly, models might incidentally learn to classify any non-viral sequences as automatically non-jelly roll sequences. For this work, this is aligned with the intended purpose, as viral JRs are the object of interest; however, in future work, it is worth investigating whether other models can find commonalities between viral and non-viral JRs.

Figure 5B shows the SJR cluster is not as well-defined (particularly at the bottom of the cluster). This is perhaps explained by the low identity among SJR sequences (Figure 3A). At the extreme top of the cluster there are mostly SJR embeddings, suggesting that for at least some sequences, embeddings alone capture SJR-like character. However, the fact that transforming these sequences to embeddings did not produce a prominent SJR cluster in general, provides evidence that SJRs may not be defined by particular sequence signatures even after embedding. In light of this, any tool which aims to predict viral SJRs would benefit from being informed by diverse sequences from many different taxa.

The VNJR cluster, on the other hand, shows a dense collection of VNJR sequences at the right of Figure 5B. Otherwise, the other VNJR sequences are also disperse. Differentiating the VNJR sequences by motif (Figure 5C) reveals a pattern. Sequences representing the β-Sheet and HK97 motifs are located almost exclusively at the VNJR cluster, but the remaining VNJR motifs are scattered throughout the ambiguous region between clusters. Taken together these results suggest sequence embeddings capture differences between some JR, most IGLBS, and VNJR sequences with Beta-Sheet and HK97 motifs, but the differences between other motifs might be more difficult to infer. This motives the need for using subsequent LR models to distinguish JR form non-JR sequences.

Lastly, Figure 5D shows misclassified sequences from the test set using the Bert-U100 LR model. There is large overlap between these misclassification results and the ambiguous group of sequences between clusters; however, there are far fewer misclassification points than total ambiguous points (Figure 5A), indicating the LR models make good use of embeddings. Notably there are a far greater number of VNJR and SJR sequences than ILBS sequences in this region, further suggesting that the Bert-U100 model uses viral sequence signatures to make classification decisions. A future examination of characteristics common to both VNJR and SJR sequences could help elucidate this phenomenon.

## Generalizability Analysis

The generalizability of the Bert-U100 model was tested on a dataset of DJR and β-strand sequences. Although the dataset was unbalanced with many more β-strand sequences than DJR sequences (ratio 25:1), this kind of imbalance mimics some real-life use cases. If the goal is filtering a small selection of “hit” sequences from a large dataset, the DJR-β sequences exemplify this use case. The results of this generalizability analysis are shown in Table 2.

**Table 2.** Bert-U100 Performance on DJR-β dataset.

|  |  |
| --- | --- |
| Accuracy | 0.92 |
| ROC AUC | 0.86 |
| False Positive Rate | 0.06 |
| False Negative Rate | 0.52 |

**Table 2.** Percent of DJRs Identified by Family.

| Family | Percent Identified JR | N | Most Common Viral Neighbor | Most Common Neighbor Motif |
| --- | --- | --- | --- | --- |
| Adenoviridae | 71 | 31 | Sinavirus | SJR |
| Ascoviridae | 0 | 11 | Secoviridae | SJR |
| Asfarviridae | 100 | 5 | Parvoviridae | SJR |
| Corticoviridae | 0 | 9 | Leviviridae | Beta Sheet |
| Faustovirus | 67 | 12 | Permutotetraviridae | SJR |
| Iridoviridae | 82 | 44 | Permutotetraviridae | SJR |
| Lavidaviridae | 67 | 9 | Microviridae | SJR |
| Marseilleviridae | 100 | 21 | Permutotetraviridae | SJR |
| Mimiviridae | 17 | 23 | Microviridae | SJR |
| Phycodnaviridae | 53 | 83 | Microviridae | SJR |
| Tectiviridae | 7 | 67 | Leviviridae | Beta Sheet |
| Turriviridae | 100 | 2 | Luteoviridae | SJR |

The Bert-U100 model exhibited a similar accuracy on the DJR-β dataset than the testing dataset (0.92 versus 0.97). However, this accuracy is bolstered by the large amount of non-DJR sequences and a low false positive rate (0.06). In other words, the model is apt ruling out non-DJR and non-JR sequences. This is particularly encouraging as the β-strand dominated subset of the DJR-β dataset is made up of only previously unseen β-strand motifs. However, there was a drop in performance regarding the false negative rate (0.52).

The model failed to label about half of all DJR folds as containing JRs. This result, though lower-than-expected, is still encouraging. All viral families in the DJR-β dataset were completely unseen during training, providing evidence of model generalizability. Moreover, the problem of identifying DJRs is distinct from identifying SJRs, meaning the current analysis might be unduly challenging for the given model. Lastly, there is evidence that adjusting the classification threshold could increase the number of DJRs identified, by still maintaining a moderate (< 0.5) false positive rate. The ROC curve in Figure 6 illustrates this point. At the current FPR of 0.06, the True Positive Rate (TPR) is approximately 0.55. However, by allowing for a FPR of approximately 0.3, the TPR increases to about 0.9, resulting in most DJR proteins being identified. Of course, deciding on a decision threshold is not always a trivial task introduces more complexity when utilizing a model. We merely remark on the threshold to show that, in principle, it is possible for the model to generalize.

**Figure 6.**
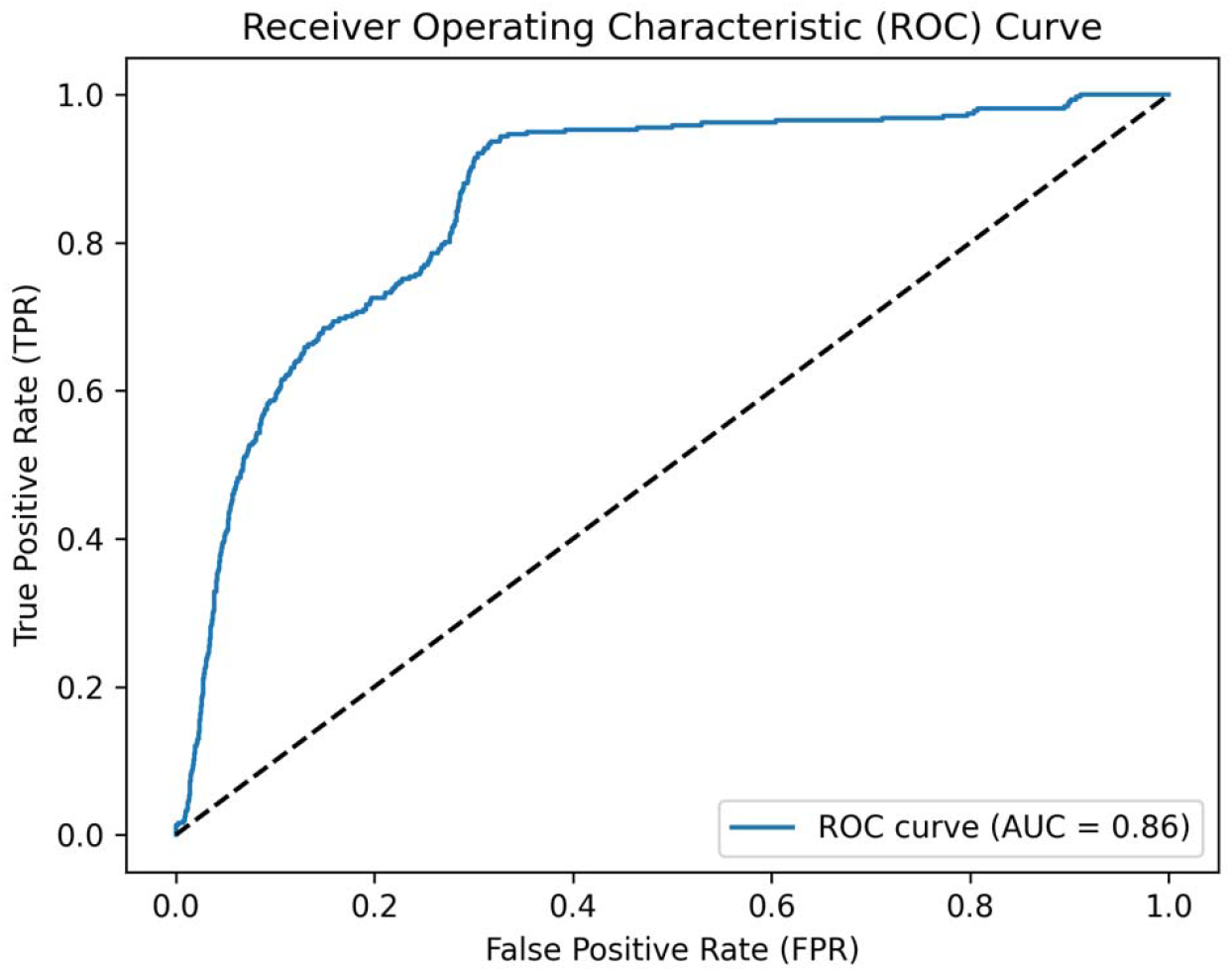
ROC curve for the generalizability dataset.

A fine-grained look into the percentage of DJR sequences illustrates performance disparities by viral family (Table 3). The data show that a large (> 66%) or a small (< 33 %) percentage of sequences were identified as JR folds per family, with only Phycodnaviridae being associated with intermediate (33%-66%) performance. Importantly, Phycodnaviridae represents 83 out of 317 entries, so the viral family has an outsized impact on the generalizability analysis. Three viral families— Asfarviridae, Marseilleviridae, and Turriviridae—were entirely identified by the model at a rate of 100%. At the other extreme, two viral families, Ascoviridae and Corticoviridae, were not identified at all by the model. For Corticoviridae this might be attributed to the family’s proximity to Leviviridae in the embedding space (Table 3). As Leviviridae has the Beta Sheet motif (non-JR), it follows that Corticoviridae would also be misclassified. Focusing on Ascoviridae, the explanation is less clear because its closest neighbor is Secoviridae—a JR virus. One possible explanation is that Ascoviridae is close but not close enough to Secoviridae in the embedding space to be recognized as a JR virus. Regardless, all these results together further illustrate that a diverse set of training sequences is required to ensure good model performance—especially for entirely unseen viral families.

## Unclassified Virus Screen

The Bert-U100 LR models were also used to predict the presence of JR folds for unclassified viruses. The results of the screen are shown in Supplemental Document S1. Two examples of novel viruses were selected to highlight the importance of studying JR viruses. Portions of their predicted capsid structures are shown in Figure 7. Both viruses exhibit the characteristic curved β-sandwich of the JR fold, and this region is predicted confidently (pIDDT > 90 for most JR residues) by AlphaFold3, corroborating the predictions of the model.

**Figure 7.**
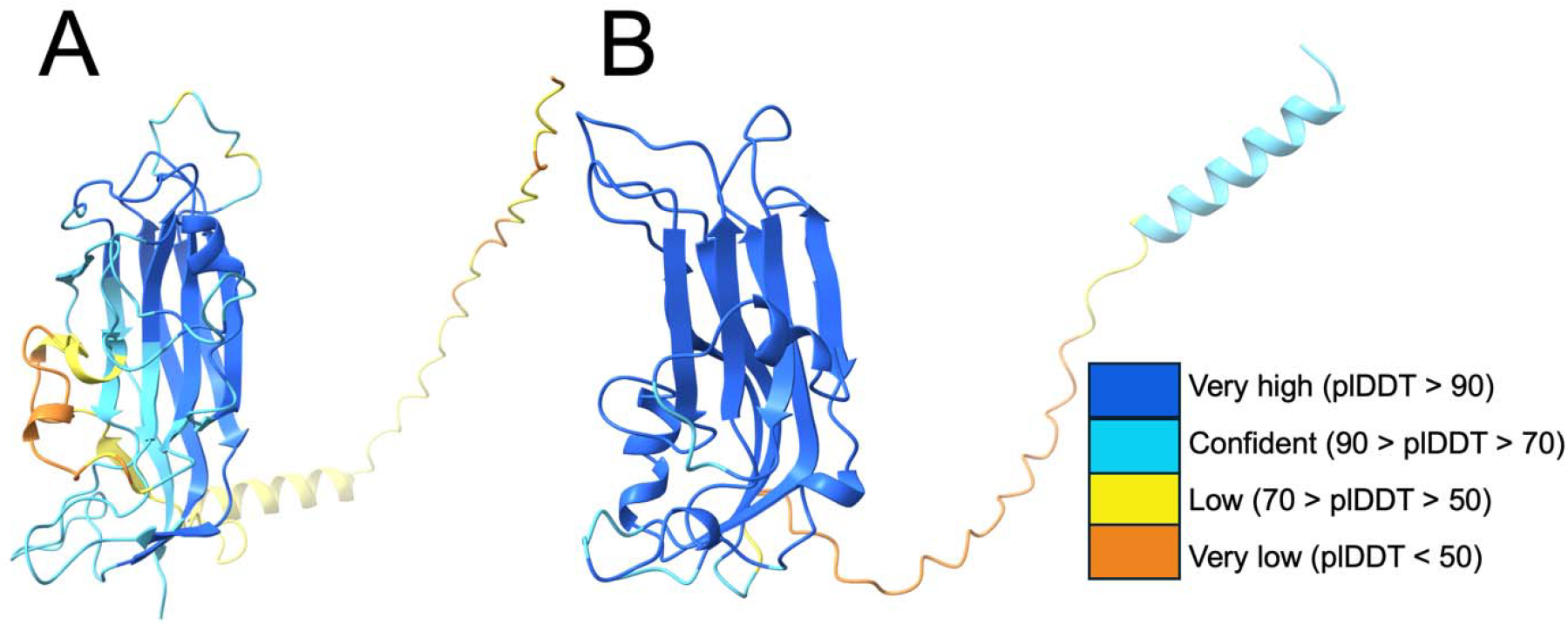
AlphaFold 3 structure predictions of unclassified JR viruses. Regions of very high, confident, low, and very low pIDDT values are shown in dark blue, teal, yellow, and orange, respectively. A) The JR portion of the Clindopec Virus 40 capsid protein was predicted to have very high or confident pIDDT values. B) All of the JR portion of the Guadeloupe mosquito virus was predicted to have very high pIDDT values.

The first of the viruses (Figure 7A) is the Clidnopec Virus 40, which is one of many novel viruses described in a study by Richard et. al.^51^ The study was geared toward characterizing the viromes of freshwater mussels (unionids) in Virginia to aid in conservation efforts. The Clidnopec virus 40 was determined to be from the Phylum Cressdnaviricota, but it lacks any subordinate taxonomical classifications. To date, all viruses within the Phylum Cressdnaviricota have been determined to bear the JR fold, and the current analysis confirms Clidnopec Virus 40 is no exception.^52,53^ Interestingly, a large percentage of all viral sequences predicted to produce JR folds comes from the Richard study. 451 viruses were identified from river sites with approximately 20% being unclassified viruses. This study highlights the potential utility of using a rapid motif-prediction tool such as JRFinder.

The second example provided is that of the Guadeloupe mosquito virus (GMV). The GMV is a viral species which has no other taxonomical information on the NCBI Taxonomy database. Present in *Aedes aegypti* and *Culex quinquefasciatus* mosquitoes, the GMV has been found in humans who present with undifferentiated Acute Febrile Illness (AFI). ^54–56^ In a study by Jitvaropas et al., blood and serum samples taken from patients in Thailand were analyzed to confirm the presence of the GMV, making sure to rule other common fever-causing pathogens endemic to the area.^54^ In total, samples from four patients were shown to contain the GMV. Although the authors mention further experimentation is required to establish a causal link between the GMV and fever, the example nevertheless serves to highlight one reason why studying JR folds is important. Other JR viruses such as the rhinovirus (family Picornaviridae), have been associated with human disease. Thus, it is probable that novel JR viruses might also be pathogenic and warrant further structural characterization in the future.

## Conclusions

Using ProtTrans LM embeddings, LR models were able to use sequences to accurately and quickly determine the presence or absence of a SJR motifs—even when challenged with diverse viral families and IGLBS sequences. Low identity among viral capsid sequences but high model performance suggests that there are similarities among viral jelly roll sequences that may be imperceptible by alignment-based methods. The performance of models appear to be embedding-independent; thus, improving JR prediction likely hinges more on data quality and diversity.

Interestingly, LM embeddings are readily able to distinguish IGLBS sequences from viral JR roll sequences as evidenced by PCA. This is likely due to the capacity of ProtTrans LMs to implicitly capture the viral or non-viral origins of sequences. PCA analysis also shows that some viral motifs appear as distinct clusters suggesting that LMs may be used to classify other types of viral motifs.

Model generalizability was tested using the Bert-U100 LR model. A challenging dataset which contained β-strand dominated motifs (non-IGLBS) and DJR folds, both of which were not seen in the training or test datasets, were used to assess generalizability. The model performed well at filtering out non-DJR sequences—as evidenced by a low false positive rate. However, many DJR sequences were predicted to be non-JR. Upon closer inspection, accurate prediction of DJR sequences as JR sequences was largely family-dependent, suggesting the model generalizes for a subset of sequences but likely requires further training to capture more diverse JR folds. Future directions involve developing other models which can more explicitly detect the presence of these DJR folds. Lastly, the Bert-U100 model was utilized to screen unclassified viruses. The capsid structures of two such viruses, the Clidnopec Virus 40 and the Guadeloupe mosquito virus, were confirmed to have JR folds by AlphaFold3. Altogether, the current work demonstrates the capacity of LLMs to make rapid predictions about viral capsid and nucleocapsid structures and motives further study of the ubiquitous JR fold.

## Methods

### Data Acquisition and Processing

Data for β-strand dominated sequences was taken from the Structural Classification of Proteins 2 (SCOP 2) database ^57,58^. Among all β-strand dominated sequences (SCOP ID: 1000001), those with the IGLBS fold (SCOP ID: 2000051) were used to form part of the balanced dataset (BD) used to assess model performance. These were chosen because they are the most numerous β-strand dominated fold and because the sandwich structure closely resembles the SJR and DJR motifs. Remaining folds were included in a generalizability dataset (GD), with the exception of sequences tagged with the jelly roll keyword. This includes JR sequences explicitly coming from viral capsid proteins (SCOP ID: 2000578) and many other non-viral JR sequences. Jelly roll sequences of viral origin were omitted from SCOP to prevent redundancy against the NCBI sequences described below. Jelly roll sequences not explicitly tagged as viral were omitted because the goal of this study is focused on viral JR prediction. The SCOP IDs of the excluded folds can be found in Supplemental Document S1. Figure 8A illustrates the subset of sequences from the SCOP database used for the BD and GD.

**Figure 8.**
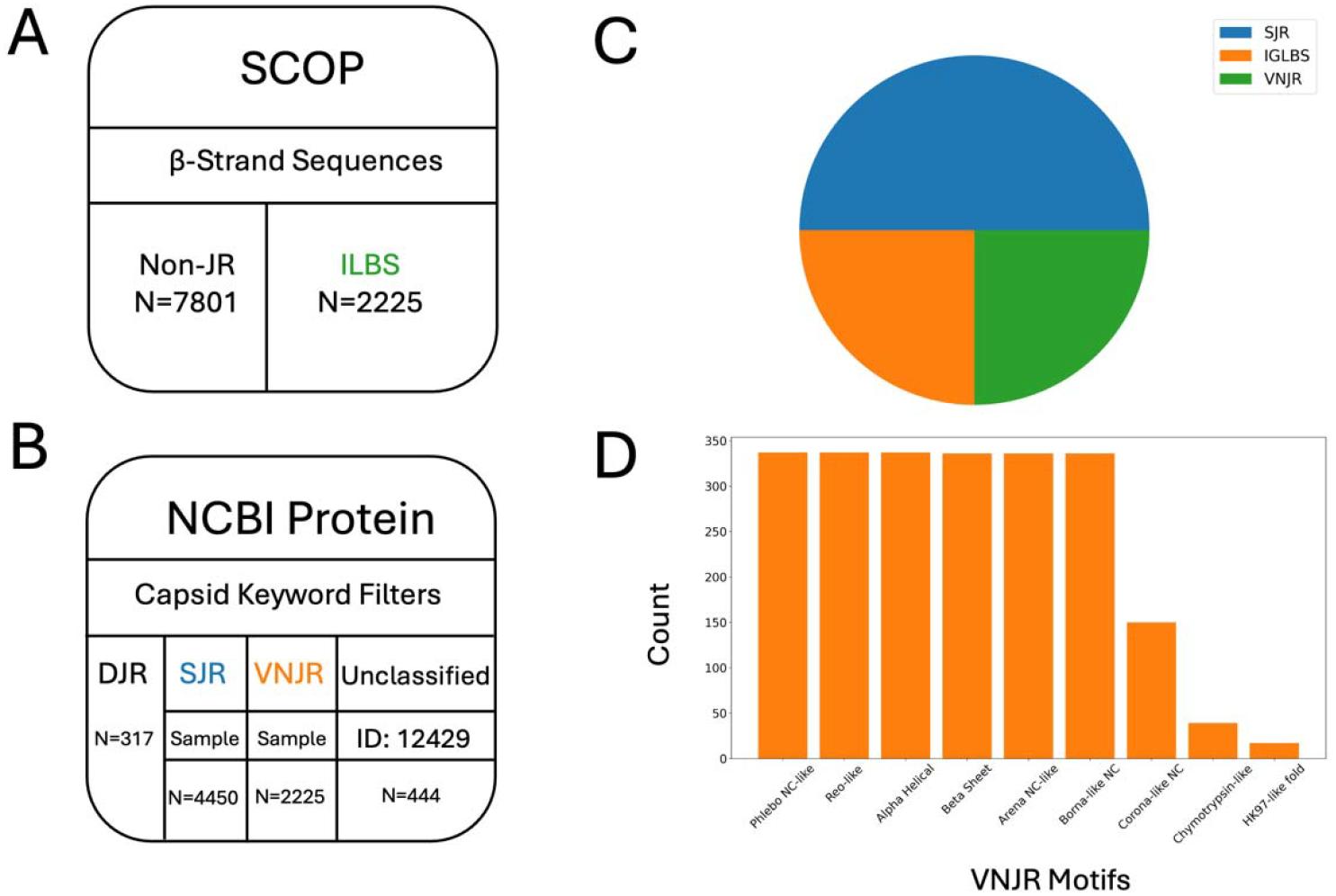
Data partitioning and composition. ILBS, SJR, and VNJR sequences are represented in green, blue, and orange, respectively. A) SCOP is the source for β-strand sequences. Among β-strand sequences, IGLBS sequences were chosen to form a part of the balanced dataset. Other non-JR sequences form a part of the generalizability dataset. B) NCBI Protein is the source for all viral sequences. All sequences were subject to keyword filters. Samples were taken from the SJR and VNJR sequences to form part of the balanced dataset. All DJR sequences were introduced into the generalizability dataset. Unclassified sequences (taxonomy ID: 12429) form the unclassified dataset. C) The balance dataset is divided such that one half of the entries represent JR sequences, and the other half represent non-JR sequences. D) A count of VNJR sequences motifs is shown. VNJR sequences were balanced by motif, when there were a sufficient number of sequences available for a given motif.

Architectural classes (structural motifs) discussed by Krupovic and Koonin were used to obtain the names of viral families associated with single jelly roll (SJR), double jelly roll (DJR), and viral non-jelly roll (VNJR) motifs.^1^ Any motif that was non-SJR or non-DJR was considered VNJR. Families with unknown or unspecified motifs were omitted. All motifs which contained the words helical, or helix were grouped into a one Alpha Helical motif. FASTA sequences associated with the resulting families were obtained from the NCBI Protein Database.^59^ Also, unclassified virus sequences (those with taxonomy ID: 12429) were obtained from NCBI.

Viral sequences were filtered using keywords present or absent in the FASTA header. Words that resulted in sequence exclusion were precursor, encapsidation, minor, partial, bridging, linking, interlacing, putative, probable, containing, and protease as these are likely associated with non-capsid and non-nucleocapsid proteins. Words required to be in the FASTA header were capsid, coat, or nucleocapsid. Of course, using this inclusion/exclusion criteria omits a number of capsid/nucleocapsid sequences and likely introduces some non-capsid/non-nucleocapsid sequences; however, such errors are difficult to avoid without pre-labelled data. Given that thousands of sequences were used for model training, a small amount of mislabeled datapoints is unlikely to significantly harm model performance. All sequences from NCBI were downloaded using the Entrez module from Biopython with the retamax parameter set to 1,000,000.^60^ Also, all aforementioned sequences (including those from SCOP) greater than 1000 amino acids were discarded. Figure 8B shows a representation of the viral sequences obtained from the NCBI Protein database.

As there were 2,225 IGLBS sequences, the same number of VNJR sequences were sampled and 5,450 SJR sequences were sampled to form the BD (Figure 8B and 8C). An attempt was made to balance VNJR motifs so that there would be equal representation of each motif in the BD (Figure 8D). The resulting BD is available in Supplemental Document S1. The balanced dataset was partitioned into training and testing datasets using scikit-learn version 1.5.2 following a 70/30 split and making sure to stratify by label.^61^ The GD dataset was made from DJR sequences and SCOP, non-JR, non-IGLBS, β-strand dominated sequences as shown in Figure 8A and 8B. The GD is available in Supplemental Document S1. Lastly, the unclassified dataset (UD), was made from unclassified viruses from the NCBI Protein database and is also available in Supplemental Document S1.

### Sequence Identity Calculations

All sequences in the balanced dataset were subject to pairwise sequence identity calculations (excluding self-self-pairs). Local sequence alignments were performed using the Biopython Align package. The Smith-Waterman algorithm and a match score of 1 were used, leaving all other parameters at their default values.^60,62^ After alignment, sequence identity was calculated for all pairs of sequences. To create average sequence identity heatmaps, all sequence identities from the relevant pairs of sequences were averaged. For example, if the Arenaviridae and Arteriviridae families were being compared, all Arenaviridae-Arteriviridae sequence identities were calculated and their scores averaged.

### Transfer Learning

A high-level view of the transfer learning process is shown in Figure 9. The first two steps of the process involve leveraging pre-trained tokenizers and transformer models from ProtTrans.^40^ These are the ProtBert-BFD100, ProtBert-UniRef100, ProtAlbert-UniRef100, ProtT5-XL-UniRef50, ProtT5-XL-BFD100, and ProtXLNet-UniRef100 transformers and their associated tokenizers. For connivence, they are renamed as Bert-BFD, Bert-U100, Albert-U100, T5-BFD, T5-U50, and XLNet-U100. All train set sequences were introduced into these tokenizers and transformers to generate embeddings. Importantly, ProtTrans embeddings are generated at the amino acid level. To obtain sequence-level embeddings, these were averaged across all amino acids in a sequence. Embeddings were calculated using PyTorch version 2.4.1 on an Ubuntu 24.04.1 environment.^63^ The motherboard used was an ASRock B550M-C with an AMD Ryzen 5 3600, 6-core, 12-thread central processing unit (CPU). Embedding generation was directed to an NVIDIA GeForce RTX 2060 graphical processing unit (GPU) using CUDA version 12.2.^64^

**Figure 9.**
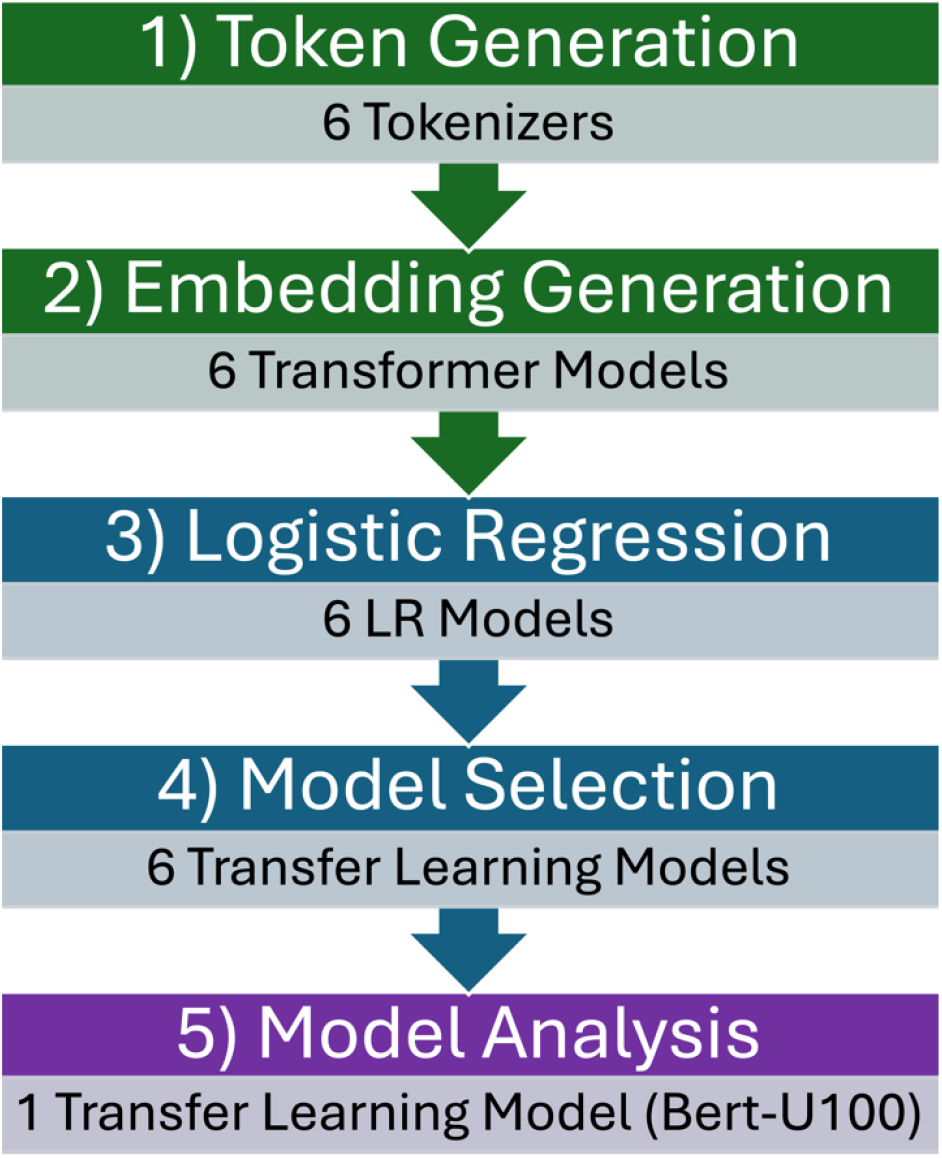
The transfer learning process. Six pre-trained tokenizers (step 1) and transformers (step 2) from ProtTrans LLMs were used to generate embeddings for input sequences. These sequences were introduced into six separate LR models (step 3). The tokenizers, transformers and LR models together represent transfer learning models. Each of the six transfer learning models was compared (step 4). One transfer learning model, Bert-U100 was chosen for its high accuracy, low false positive rate, and quick embedding generation, and subjected to subsequent analysis (step 5).

Train set embeddings were scaled to zero mean and unit variance using the scikit-learn StandardScaler and the scaled embeddings were subsequently used to train six LR models (step 3). LR models were trained using PyTorch (one linear layer and one sigmoid activation function). The loss function was binary cross entropy. The optimizer used was the Adam optimizer with a learning rate of 0.0001. Each LR model was trained for 1,000 epochs. The batch size was equal to the number of entries in the train set. For each epoch, accuracy was calculated on the train set. The model parameters which produced the highest accuracy were saved. If an identical accuracy was calculated at a later epoch, the parameters of the earlier epoch were saved. During training, test set sequences were also tokenized, subjected to embedding and scaled (using the saved train set scalers) and the binary cross entropy was calculated for these. Learning curves demonstrate a plateau in both train and test set cross entropy losses, indicating models had not experienced overfitting during the 1,000 training epochs. Learning curves are available in Supplemental Document S2.

Following training, the models were evaluated on the scaled test set embeddings (step 4). The performance metrics for all models were calculated using the following equations, where TP, TN, FP, and FN are the number of true positive, true negative, false positive, and false negative predictions in the test set:

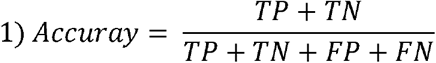

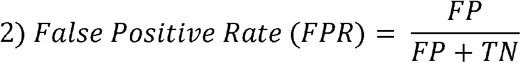

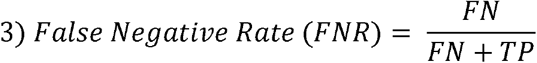

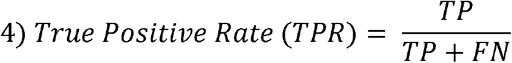

The model selected for subsequent analysis was Bert-U100 (step 5). Principal component analysis of the Bert-U100 embeddings (unscaled) was conducted using scikit-learn. The generalizability analysis of the Bert-U100 model was done using the GD described above. The most common neighbor for each DJR family (Table 2) was determined by comparing DJR to SJR or VNJR sequences. For each DJR sequence, the nearest SJR or VNJR sequence in the embedding space (determined by Euclidean distance) was found. The family of the nearest SJR or VNJR neighbor for each DJR sequences was recorded, and results were grouped for each DJR family. The most common (mode) SJR or VNJR family for each DJR family was reported. Lastly, the unclassified virus screen was conducted using the UD.

### Figure Generation

Matplotlib version 3.9.2 was used for figure generation.^65^ Seaborn was used to create heatmaps.^66^ The ROC curves were generated using the scikit-learn. Operations which required manipulating tabular data were completed using pandas version 2.2.3.^67,68^ Manipulation of matrix data was done with NumPy version 2.1.1.^69^ USCF ChimeraX version 1.9 was used for illustrating the differences between the motifs in the balanced dataset.^70^ Predicted JR capsid structures were generated using AlphaFold3.^33^

## Acknowledgements

Research reported in this publication was supported by the Welch Foundation under Grant Number AH-2126-20220331. LL and XC received support from National Institute of General Medical Sciences under award numbers SC1GM132043 and R01GM129525, respectively. LL and XC also received support from National Institutes on Minority Health and Health Disparities (NIMHD) under award number U54MD007592. The content is solely the responsibility of the authors and does not necessarily represent the official views of the funding agencies.

