## Supplemental File 2 for "JRSeek: Artificial Intelligence Meets Jelly Roll Fold Classification in Viruses"

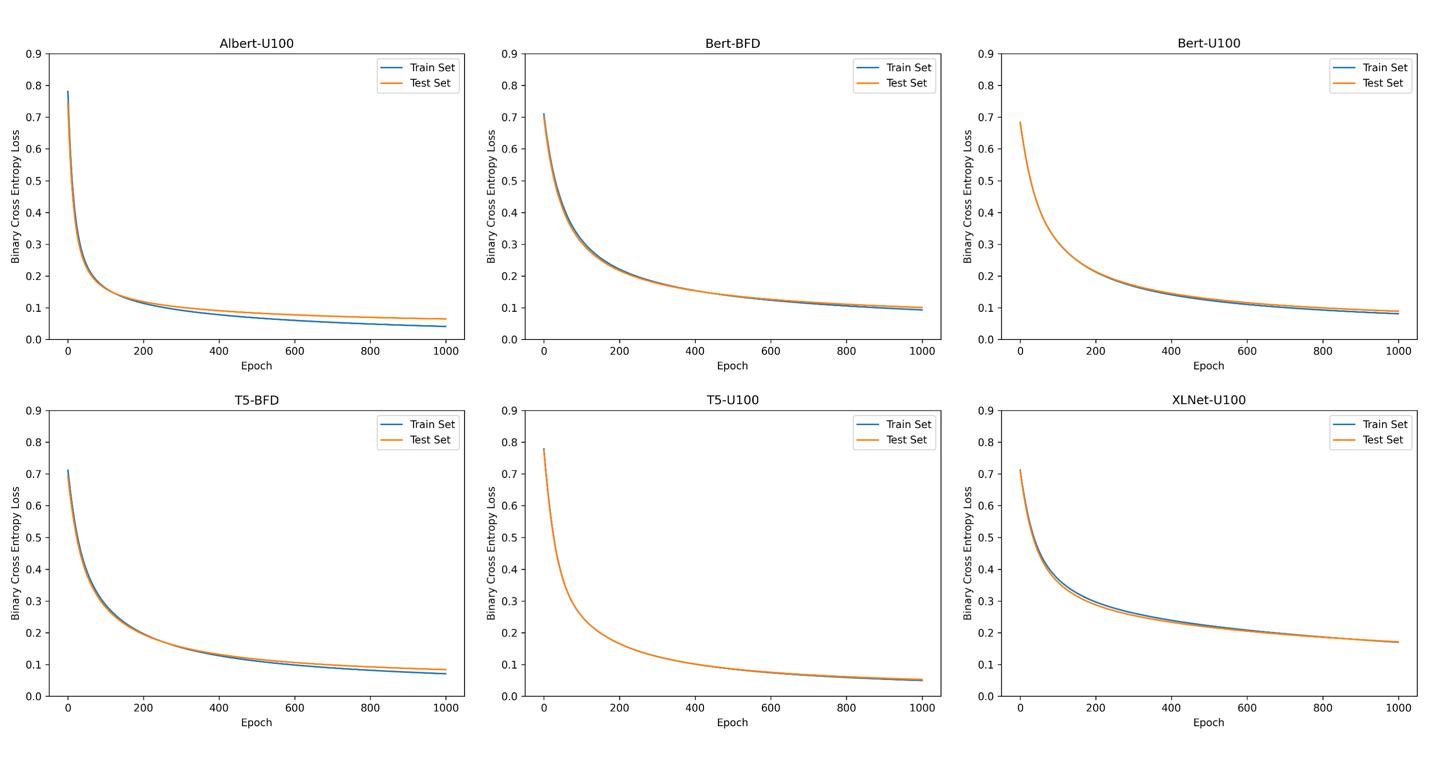

Figure S2.1. Learning curves for the six logistic regression models. The titles represent the embeddings used.
